# Th2-like T-follicular helper cells promote functional antibody production during *Plasmodium falciparum* infection

**DOI:** 10.1101/2020.05.18.101048

**Authors:** Jo-Anne Chan, Fabian de Labastida Rivera, Jessica Loughland, Jessica A Engel, Hyun Jae Lee, Arya SheelaNair, Bruce D Wines, Fiona H Amante, Lachlan Webb, Pamela Mukhopadhyay, Ann-Marie Patch, P. Mark Hogarth, James G Beeson, James S McCarthy, Ashraful Haque, Christian R Engwerda, Michelle J Boyle

## Abstract

The most advanced malaria vaccine only has approximately 30% efficacy in target populations, and avenues to improve next generation vaccines need to be identified. Functional antibodies are key effectors of both vaccine induced and naturally acquired immunity, with induction driven by T-follicular helper cells (TfH) CD4+ T cells. We assessed circulating TfH (cTfH) responses and functional antibody production in human volunteers experimentally infected with *Plasmodium falciparum*. Longitudinal single-cell RNA-sequencing of cTfH revealed peak transcriptional activation and clonal expansion of major cTfH subsets occurred at day 8 following infection and a population structure of cTfH capturing phenotypical subsets of Th1- and Th2-like cells. Among 40 volunteers, infection resulted in the emergence of activated ICOS+ cTfH cells. During peak infection, activation was restricted to Th2-like cTfH cells, while Th1-like cTfH cell activation occurred one week after treatment. To link cTfH activation to antibody induction, we assessed the magnitude and function of anti-malarial IgM and IgG after infection. The functional breadth and magnitude of parasite-specific antibodies was positively associated with Th2-cTfH activation. In contrast, Th1-cTfH activation was associated with the induction of plasma cells, which we have previously shown have a detrimental role in germinal cell formation and antibody development. Thus, we identified that during *P. falciparum* malaria infection in humans, the activation of Th2-cTfH but not other subsets correlates with the development of functional antibodies required for protective immunity. Data for the first time identify a specific cellular response that can be targeted by future malaria vaccines to improve antibody induction.

## Introduction

*Plasmodium falciparum* malaria remains one of the most important infectious diseases globally, with over 200 million clinical cases and up to 500,000 deaths annually *(1)*. Antibodies play a major role in naturally-acquired and vaccine-induced protective immunity to malaria, but factors determining their optimal induction and maintenance over time is poorly understood. The first licensed malaria vaccine is RTS,S, which induces antibodies targeting sporozoites to prevent infection *(2–4)*, and large scale Phase IV implementation trials have recently commenced in parts of Africa. However, RTS,S has only moderate efficacy and short-lived protection in infants and young children *(2)*. Thus, understanding the induction and maintenance of antibodies is crucial for developing vaccines with greater efficacy and durability.

Antibody production, both during infection and vaccination, is driven by CD4^+^ T-follicular helper (TfH) cells which promote the development of memory and class-switched, affinity-mature antibody-producing B cells *(5)*. Due to the central importance of TfH cells in antibody induction, optimized targeting of these cells has been proposed as an avenue for improving vaccines *(6, 7)*. Indeed, recent studies of the role of differential dosing and vaccination schedule in the efficacy of RTS,S, suggest that induction of vaccine specific TfH is one key determinant of protective efficacy *(8)*. To develop strategies for targeting TfH cells in malaria vaccines, improved understanding of the role of this key CD4^+^ T cell subset in helping B cells produce protective antibodies during malaria is required.

TfH cells primarily function within germinal centres (GC) *(5)*. However, in humans, subsets of peripheral blood circulating CD4^+^ T cells have been reported to express CXCR5 and programmed death-1 (PD1) molecules, and to share phenotypic, functional and transcriptional profiles of lymphoid TfH *(9, 10)*. In further studies, significant clonal overlap between peripheral and tonsillar GC TfH has been described *(11, 12)*. Upon activation, circulating TfH (cTfH) cells express the co-stimulatory marker ICOS, which is crucial for development and function of lymphoid TfH cells *(13–15)*, as well as other activation markers including CD38 and Ki67 *(5, 16)*. ICOS+ cTfH have been associated with vaccine induced antibody induction *(16–21)*. By classifying CXCR3 and CCR6 expression cTfH can be segregated into three different T helper (Th)-like subsets: Th1-(CXCR3+CCR6-), Th2-(CXCR3-CCR6-) and Th17-(CXCR3-CCR6+). Th2-cTfH cells exhibit transcriptional profiles most closely related to GC TfH cells *(9)* and, along with Th17-cTfH cells, have the greatest capacity to activate naïve B cells *in vitro (10)*. In contrast, Th1-cTfH cells appear to have important roles in the activation of memory B cells *(19)*. The relative importance of specific cTfH cell subsets in driving antibody responses appears to be influenced by the type of pathogen and context of the exposure. For example, Th2-cTfH cells have been associated with the acquisition of broadly neutralizing antibodies against HIV *(9)*, while Th1-cTfH cells are linked to antibodies following influenza and HIV vaccination *(19, 22, 23)*. cTfH cell activation during *Plasmodium* infection in humans have only been investigated in a few studies *(24, 25)* and to date no studies have linked a specific cTfH cell subset anti-malaria antibody production and functional activity.

Previous studies have indicated that *P. falciparum* infection results in the preferential activation of Th1-cTfH cells. In Malian children with acute symptomatic disease, only Th1-cTfH cells were activated. However, activation of Th1-cTfH cells did not correlated with total anti-parasitic IgG responses *(24)*. Further, Th1-cTfH cells have been linked to the development of atypical memory B cells *(26)*, which have exhausted phenotypes *(27, 28)*. It has also been hypothesised that Th1-TfH cells may preferentially promote short-lived B cell responses, including plasmablasts *(29)*, which we recently reported to act as a metabolic ‘sink’ constraining GC reactions, to the detriment of antibody and memory B cell responses *(30)*. Further supporting a negative role of Th1-cTfH cells in malaria, co-administration of viral vectored vaccines with RTS,S skewed the response towards a Th1-cTfH cell activation phenotype which hindered development of functional antibodies and reduced vaccine efficacy *(31)*.

We hypothesized that Th2-cTfH, rather than Th1-cTfH, cell activation during *P. falciparum* infection is required to promote effective anti-malarial antibody production. To test this hypothesis, we employed experimental human volunteer infection studies using the induced blood stage malaria (IBSM) system *(32)*). We first analysed cTfH cell population dynamics via single-cell RNA sequencing (scRNA-seq), and then quantified cTfH cell activation and antibody magnitude, type, and functional activity during *P. falciparum* infection in a group of 40 participants. Our results characterise, for the first time, cTfH cell subsets and phenotypes that were associated with production of functional antibodies in human malaria.

## Results

### Single-cell transcriptomics of cTfH during induced blood stage malaria

To assess cTfH cell population structure and activation dynamics during infection, first we sorted CXCR5^+^ CD45RA^−^ CD4^+^ T cells from one study participant prior to, during, and following IBSM (day 0, peak infection/day 8, day 15 and day 36) and analyzed cells via droplet-based scRNA-seq. After filtering for poor quality transcriptomes, 14,275 cells were selected for downstream transcriptome analysis (day 0: 4777; day 8: 3565, day 15: 2978, day 36: EOS, 2955). Similar numbers of genes (1314±339, mean ± S.D.) and unique molecular identifiers (UMIs, 4743±1636) were detected at each time points. Data were integrated across the four timepoints *(33, 34)*, and dimensionality reduction and 2D-embedding performed via UMAP. Nine clusters were identified, ranging in prevalence from ~50% for cluster 1 to <0.5% for clusters 8 & 9 (Figure 1A). Clusters were present at a similar proportions across all time points suggesting a stable population structure during and following infection (Figure 1B).

**Figure 1:**
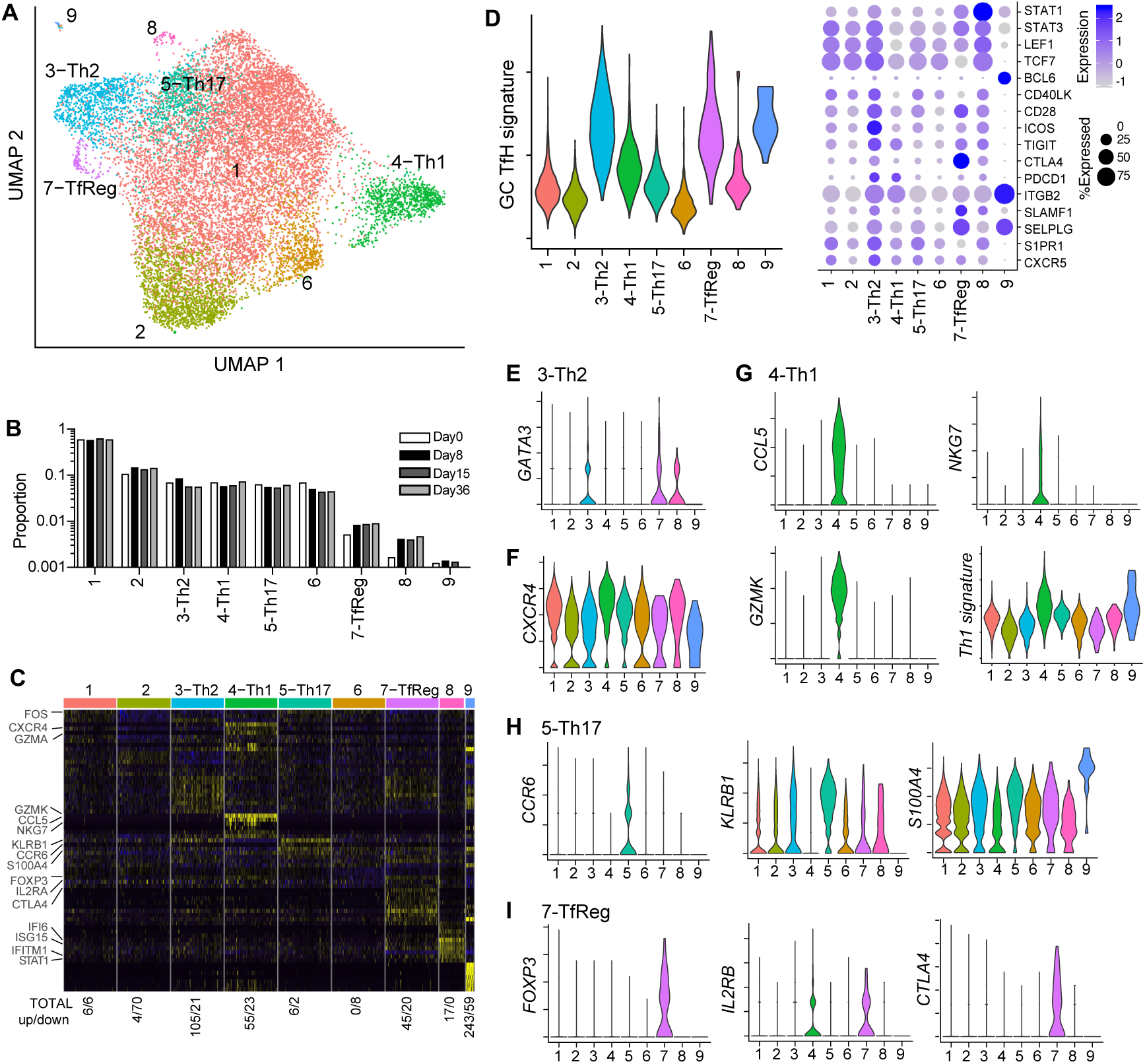
Analysis of TfH subsets by single cell RNA sequencing. **A)** CXCR5+CD45RA-CD4 T cells were sorted at day 0, 8, 15 and 36 following P. falciparum infection in one study participant and analysed by scRNA-seq. UMAP plot following integration of data sets across time points. Cells are coloured by cluster. **B)** Contribution of cells from each infection time point in each cluster as a proportion of total cells. **C)** Top 10 conserved defining genes in each cluster (independent of infection time point). Complete list of defining differential expressed genes can be found in Supplementary Table 1. Total number of significantly differentially expressed genes (p<0.01) in each cluster is indicated below (total up/down). **D)** GC TfH conical markers across clusters and GC TfH signature genes calculated as a combined expression of GSE50391, CXCR5hiCD45RO verses CXCR5negCD45ROneg tonsil samples. **E-I)** Violin plots show single-cell expression of indicated transcripts across identified clusters. For Th1-signature expression is total combined levels of Th1-signature genes (36). See also Supplementary Figure S1 and Supplementary Table S1.

To identify clusters most similar to cTfH cells amongst the CXCR5^+^ CD4^+^ T cell population, common cluster marker genes were identified *(33, 34)* (Figure 1C, **Supplementary Table S1**), and clusters were examined for GC TfH cell gene signatures *(9)* (Figure 1D). Cluster 3 had the highest overall GC TfH cell signature, and also high expression of the Th2-master transcription factor *Gata3* (Figure 1D/E), consistent with being a Th2-cTfH population *(9)*. Cluster 3 also exhibited lower expression of *Cxcr4*, which influences positioning within GC *(35)* (Figure 1F, **Supplementary Table 1**). Relatively lower CXCR4 expression within Th2-cTfH cells compared to other subsets was confirmed by flow cytometry in additional healthy donors (**Supplementary Figure S1A**). Cluster 4 expressed high levels of Th1-associated genes *Ccl5*, *Nkg7* and *GzmB*, and a high overall Th1-signature *(36)*, suggesting Cluster 4 as Th1-cTfh cells (Figure 1G). The restriction of granzyme B to Th1-cTfH cells was confirmed by flow cytometry (**Supplementary Figure 1B**). Cluster 5 uniquely expressed *Ccr6*, the marker previously employed to identify Th17-cTfH cells, as well as *Klrb1* (encoding CD161) and *S100A4* (Figure 1H). Cluster 7 exhibited both a strong GC TfH cell signature (Figure 1D) and unique high expression of *Foxp3, Il2ra* and *Ctla4*, consistent with a TfH-regulatory (TfReg) profile *(37)* (Figure 1I). Cluster 7 also shared transcriptional similarities with Th2-like Cluster 3 (Figure 1C), and flow cytometry confirmed TfReg cells were predominately Th2-like (**Supplementary Figure 1C**). Of the remaining clusters, cluster 8 was defined by high expression of Type I IFN-associated genes, including *Stat1, Ifitm1, Isg15*, and *If16* (Figure 1C), while the frequency of Cluster 9 (~0.1%, <50 cells total) was too low to be analyzed. Clusters 1, 2 and 6 had lower expression of GC signature genes and expressed no clear features of Th1-, Th2-, Th17-cTfH or TfReg cells (Figure 1C/D). Cluster 1 and 6 had only limited marker genes, and Cluster 2 was unique only in its down-regulation of 70 out of 74 marker genes, including *Fos* and *Jun* (Figure 1C, **Supplementary Table 1**). Thus, unbiased transcriptomic assessment of gene transcription in circulating CXCR5^+^ CD4^+^ T-cells revealed a population structure for cTfH cells consistent with reported flow cytometric sub-setting using CXCR3 and CCR6 which delineates cTfH into Th2-, Th1-, Th17-like populations.

### cTfH are activated and clonally expanded during peak infection

To assess cTfH cell activation across subsets and timing of activation during volunteer infection, we compared transcription profiles between day 0 and all post-infection time points. Across all cells, 75 genes were differentially-expressed at day 8 compared to day 0, which contracted to 10 and 28 genes at day 15 and 36 respectively. This pattern was similar amongst Clusters 1-6, while Clusters 7-9 appeared not to be activated during infection (Figure 2A). Th2-like and Th1-like Clusters 3 and 4, respectively, had the highest number of upregulated genes at day 8 compared to day 0. Upregulated genes included those associated with TfH cell development *Stat1 and Tcf7*, along with Type I IFN-associated genes *(38)*, *Ifitm1*, *Ifitm2*, *Ifitm3, Irf1, Tap1, Ly6e, Gbp1/2* and *Stat1* (Figure 2B). A Type I IFN gene signature was up-regulated across Clusters 1-6, with the greatest upregulation in Th2-like Cluster 3 (Figure 2C). By day 15 and 36 post-infection, Th2-like Cluster 3 appeared to maintain transcriptional changes for longer than other Clusters, and included a marked down regulation of CXCR4 at day 15, which was also down-regulated in other Clusters at day 36 (Figure 2A/B).

**Figure 2:**
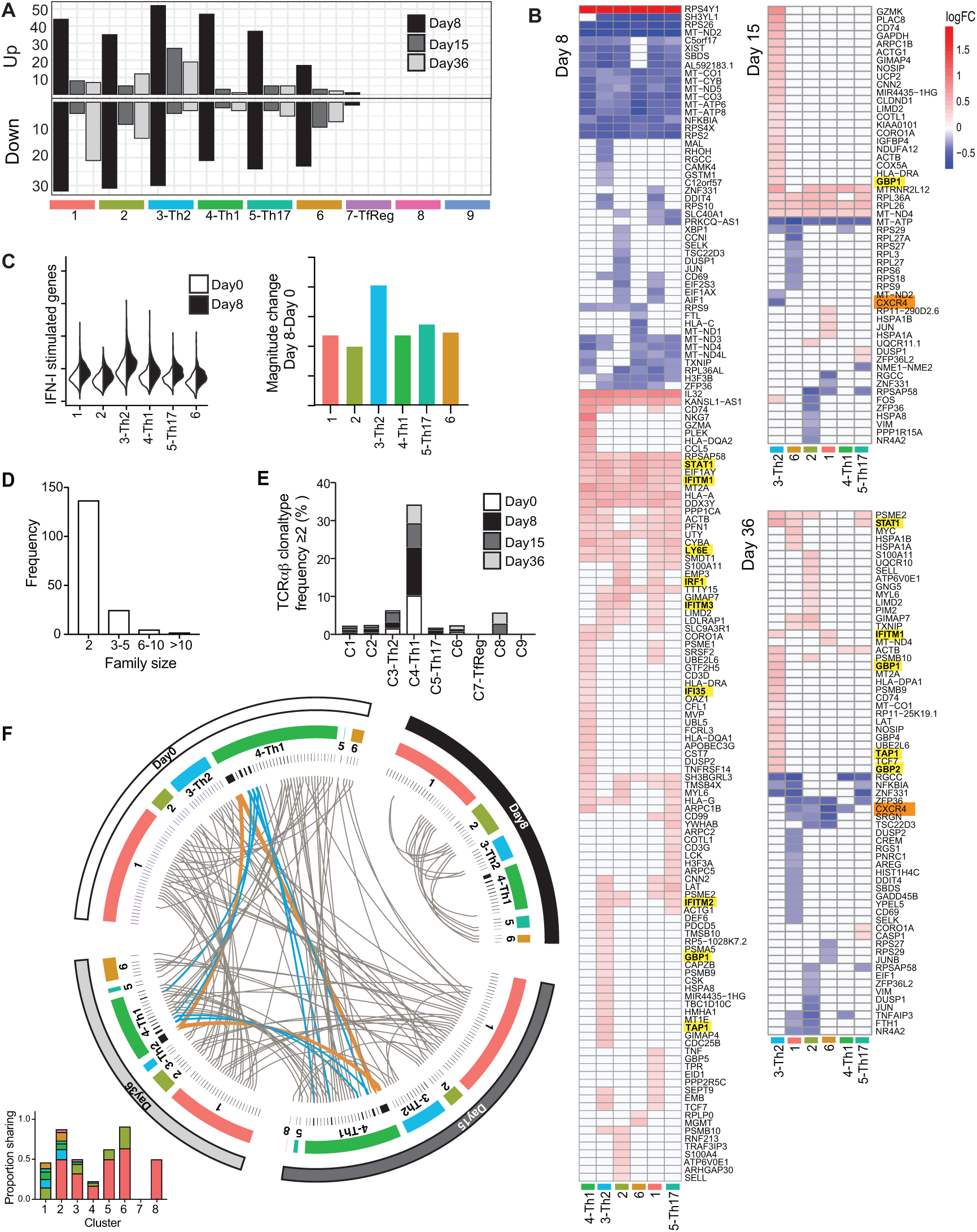
Transcriptional changes and clonal relationships within TfH cells following infection. **A)** Differentially expressed genes at day 8, 15, 36 compared to day 0 timepoints were identified for each TfH cell cluster. Numbers of up and down regulated genes at each timepoint following infection within each cell cluster. **B)** Log Fold Change (FC) of differentially expressed genes in clusters C1-C6 at each time point. Genes and clusters were grouped by euclidean clustering. Genes associated with Type I IFN signaling are highlighted in yellow, and Cxcr4 is indicated in orange. **C)** Overall accumulative expression of genes identified as type 1 IFN response genes in CD4 T cells (38), at day 0 and day 8 within clusters C1-C6 (left panel). Magnitude change of Type I IFN responses genes (right panel). **D**) Cells with shared TCRαβ sequences were considered clones. Bar graph of frequency of clonal families with family sizes 2, 3-5, 6-10 and 10+ cells. **E)** Bar graph of the percentage of cells from cluster that were clonally expanded, having at TCRαβ chain with a frequency ≥2. **F)** Circos diagram of relationship between clones across time points and clusters. Clones are grouped by day and by Cluster. Ribbons connect sister clones. Grey ribbons connect clonal size 2, blue ribbons connect clonal sizes 3-10 and orange connect clonal sizes greater than 10. Bar graph insert shows the proportion of all clones within each cluster shared in with other clusters.

We also assessed clonal expansion in cTfH cells during IBSM. Paired TCR and TCR sequences were captured in 9980/14,275 cells (69.9%). Of these, 473 (4.7%) cells were clonally expanded, sharing TCRαβ with at least one other cell. 172 unique clones were identified and family sizes ranged from 2-35 (Figure 2D). TCRαβ were linked to Cluster and timepoint annotation. Clonal expansion was most frequently in Th1-like Cluster 4 (34.3%), and Th2-like Cluster 3 (6.45%), with clonal expansion present before, during and after infection (Figure 2E). Clones were shared both between days and across Clusters (Figure 2F). However, all clones present at day 8 were unique to that timepoint, suggesting displacement of existing clonal cTfH cells by expansion of parasite-specific cTfH cells during peak infection. Clone sharing between Clusters was evident, particularly of clones within Cluster 1 and Cluster 2. In contrast, greater than 50% of clones in Th2-like Cluster 3 and Th1-like Cluster 4 were unique to those clusters, and only one clone was shared between Th1-like Cluster 4 and Th2-like Cluster 3, consistent with previous data showing low clonal overlap between CXCR3^−^ and CXCR3^+^ cTfH cells *(12)*. Taken together, longitudinal scRNAseq analysis of a single IBSM participant suggested that cTfH cell subsets, including Th2-like cells, had been differentially activated and clonally expanded, and that week 2-3 of infection was an appropriate time period for assessing cTfH cell activation in this experimental human model of malaria.

### cTfH cells upregulate multiple activation markers during induced blood stage malaria

Having characterised cTfH cell dynamics in one individual by scRNAseq, we next quantified cTfH cell activation in a cohort of IBSM participants (n=40), on day 0 (prior to inoculation), day 8 (peak infection/treatment time point), day 14/15 and end-of-study (EOS) (**Supplementary Figure S2**). Activation of cTfH cells was assessed using commonly used activation markers ICOS and CD38. While the frequency of cTfH cells amongst circulating CD4^+^ T cells remained unchanged (Figure 3A/B), ICOS was up-regulated at peak infection, and further increased by day 14/15 and EOS (Figure 3C/D). There was no correlation between pre-infection levels of ICOS and the magnitude of upregulation during infection (**Supplementary Figure S3A**). Furthermore, ICOS-upregulation was not correlated with parasite biomass which varied substantially amongst participants (**Supplementary Figure S3B/C**). ICOS^+^ cTfH cells, but not ICOS^−^ counterparts, also expressed the activation marker CD38, which increased following treatment at day 14/15 (Figure 3E/F). Ki67 and HLA-DR expression was also evident amongst cTfH and was higher on ICOS^+^, compared to ICOS^−^ cTfH cells (**Supplementary Figure S4**). This indicated that across a cohort of participants, and consistent with our scRNAseq analysis of one individual, that IBSM induced cTfH cell activation.

**Figure 3:**
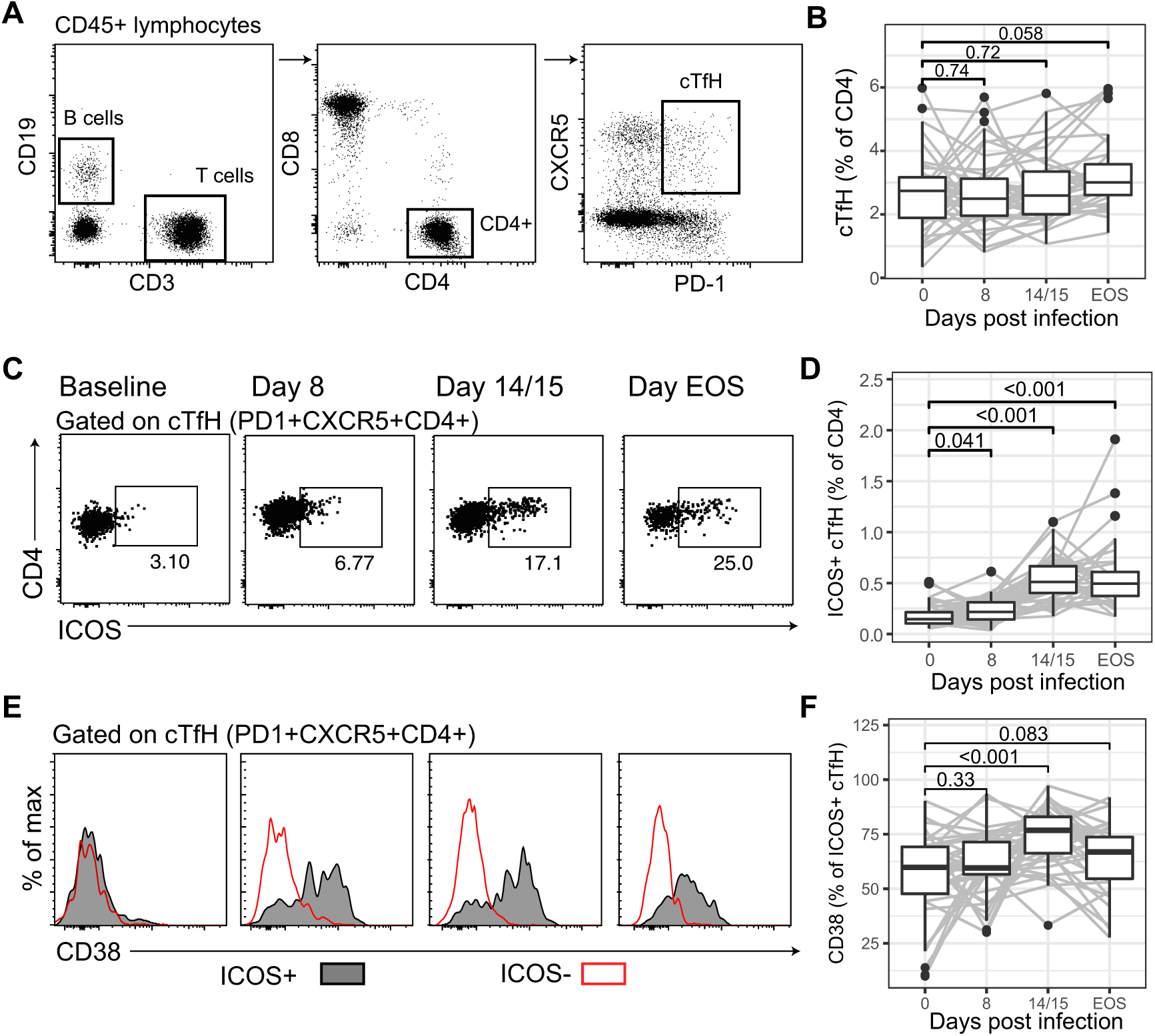
cTfH are activated in P. falciparum infection. **A)** cTfH (PD1+CXCR5+CD4) T cells were assessed by flow cytometry at day 0, 8, 14/15 and EOS. For full gating strategy see Supplementary Figure S2. **B)** cTfH cells as a frequency of total CD4 T cells after infection. **C)** Representative fluorescence-activated cell sorting (FACs) of activated (ICOS+) cTfH cells following infection. **D)** Activated cTfH (ICOS+) cells as a frequency of CD4 T cells. **E)** Representative FACs plots of CD38 expression on activated (ICOS+) and non-activated (ICOS-) cTfH cells. **F)** CD38 positivity as a frequency of activated ICOS+ cTfH cells. For **B/D/F** Paired Wilcoxon signed rank test compared to day 0 are indicated. See also Supplementary Figure S2-S4.

To identify specific cTfH cell subsets activated during IBSM, we examined cTfH as Th1-, Th2- and Th17-subsets based on CXCR3 and CCR6 expression (**Supplementary Figure S2**). At day 8 (peak infection), the proportion of cTfH cells identified as Th2-like increased compared to day 0, whereas Th1-cTfH proportions decreased. After drug treatment, the proportion of Th2-cTfH returned to baseline, whereas there was a marked increase in Th1-cTfH proportions (Figure 4A). Similarly, activated ICOS^+^ Th2-cTfH frequencies increased at peak infection, while ICOS^+^ Th1-cTfH was low at day 8 and dramatically increased after drug treatment (Figure 4B). While significant changes were detected among subjects, it should be noted that there was substantial variation in the subset composition of cTfH cell activation between individuals (Figure 4C). Together, these data indicated that cTfH cell subsets were activated with different kinetics during IBSM; Th2-cTfH cells were predominately activated early during peak infection, while Th1-cTfH cell activation occurred post-treatment, and subsequently dominated the cTfH cell response.

**Figure 4:**
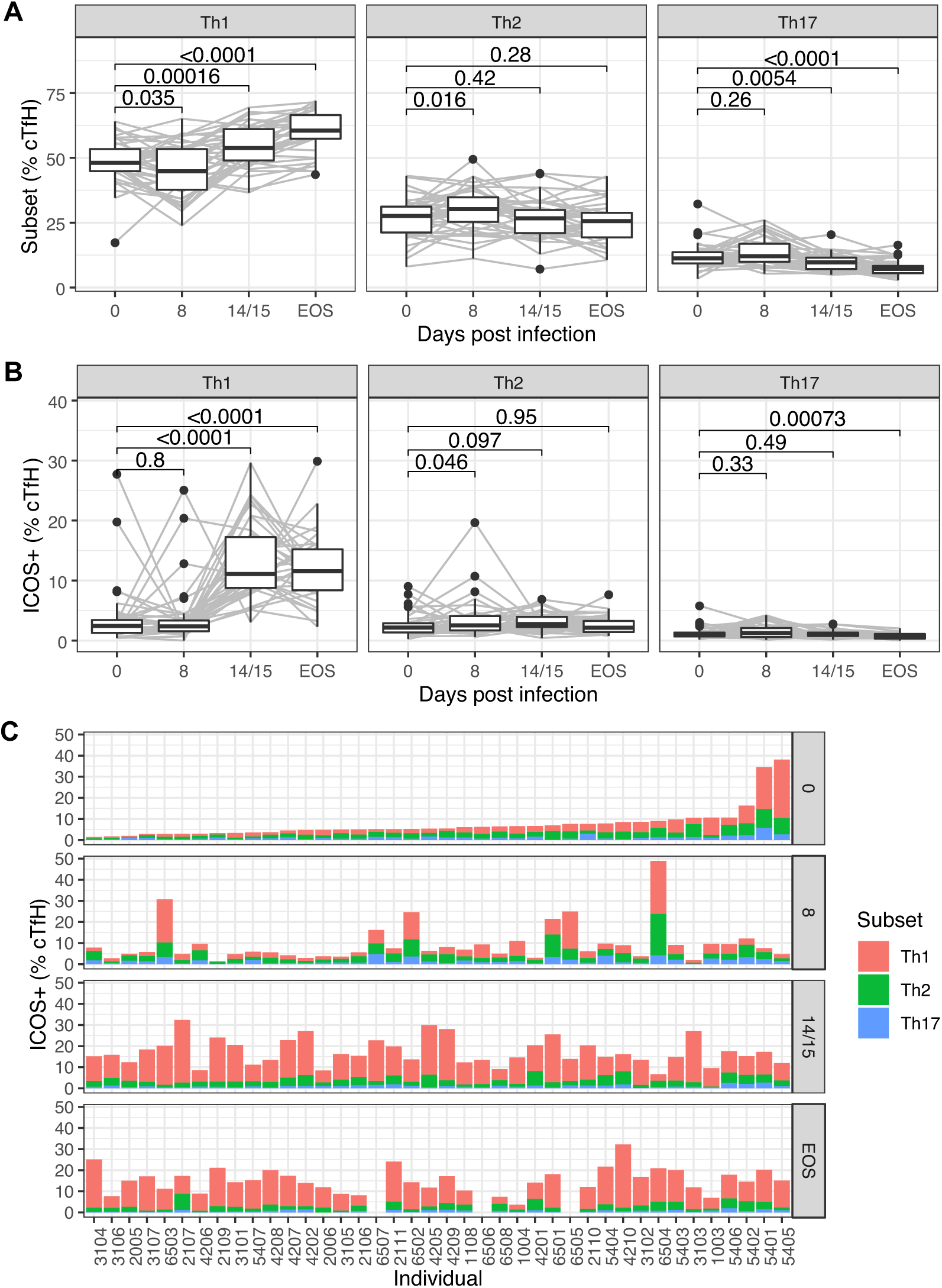
Th2-cTfH are activated early in P. falciparum infection. cTfH (PD1+CXCR5+ CD4) cells were differentiated into subsets based on CXCR3 and CCR6 expression into Th1-(CXCR3+CCR6-), Th2-(CXCR3+CCR6+) and Th17-like (CXCR3-CCR6+) cell subsets. **A)** Subsets as a frequency of cTfH cells after infection. **B)** Activated ICOS+ subsets as a frequency of cTfH cells after infection. **C)** Activated ICOS+ subsets as a frequency of cTfH cells for each individual study participant following IBSM. Participants are ranked based on the frequency of ICOS+ cTfH cells prior to infection. Note – for participants 6507, 6505 and 6506 no sample was available at end-of-study time-point (NA). For **A/B** Wilcoxon pair t-test to day 0 are indicated.

### Isotype-switched and functional antibodies are induced following IBSM

To examine the relationship between cTfH cell activation and the development of protective humoral immunity, we next analysed parasite-specific antibodies generated during IBSM. We quantified IgM and IgG subclass *(39)*, along with the magnitude of functional antibodies that fixed complement factor C1q *(40, 41)*, which is the first step in classical complement activation, bind Fc_γ_-receptors (Fc_γ_R) that are involved in opsonic phagocytosis and antibody dependent cellular cytotoxicity [36,37], and mediate opsonic phagocytosis (OPA) by THP-1 monocytes *(42)*. We measured responses to intact merozoites *(43)*, the parasite stage that invades RBCs, and to the abundant and immunodominant merozoite antigen and vaccine candidate, MSP2 *(44–47)*. Following IBSM, IgM and all IgG subclasses recognising merozoites and recombinant MSP2 were increased at EOS (Figure 5A/B). The proportion of individuals responding was highest for IgM and IgG1. Similarly, all functional antibody responses were significantly increased after infection except for antibody promoting Fc_γ_RIIa binding to merozoites (Figure 5C/D). The prevalence of functional antibodies was highest for antibodies that crosslinked Fc_γ_RIIIa to merozoites and MSP2. Functional responses (Fc_γ_R binding, C1q fixation and OPA) were moderately, and significantly, correlated with IgG1 (R=0.49-0.76), and less commonly with IgG3 (R=0.32-0.46). However, each functional response was only moderately correlated to other functions, indicating that functional responses were not equally co-acquired (Figure 5E). To understand the overall breadth and composition of an individuals’ response, each antibody response was categorized as negative (below positive threshold) or low/high (based on below/above median value of all positive responses). The overall composition of induced antibody responses between individuals was diverse, however there was a clear hierarchy of antibody acquisition (Figure 5F).

**Figure 5:**
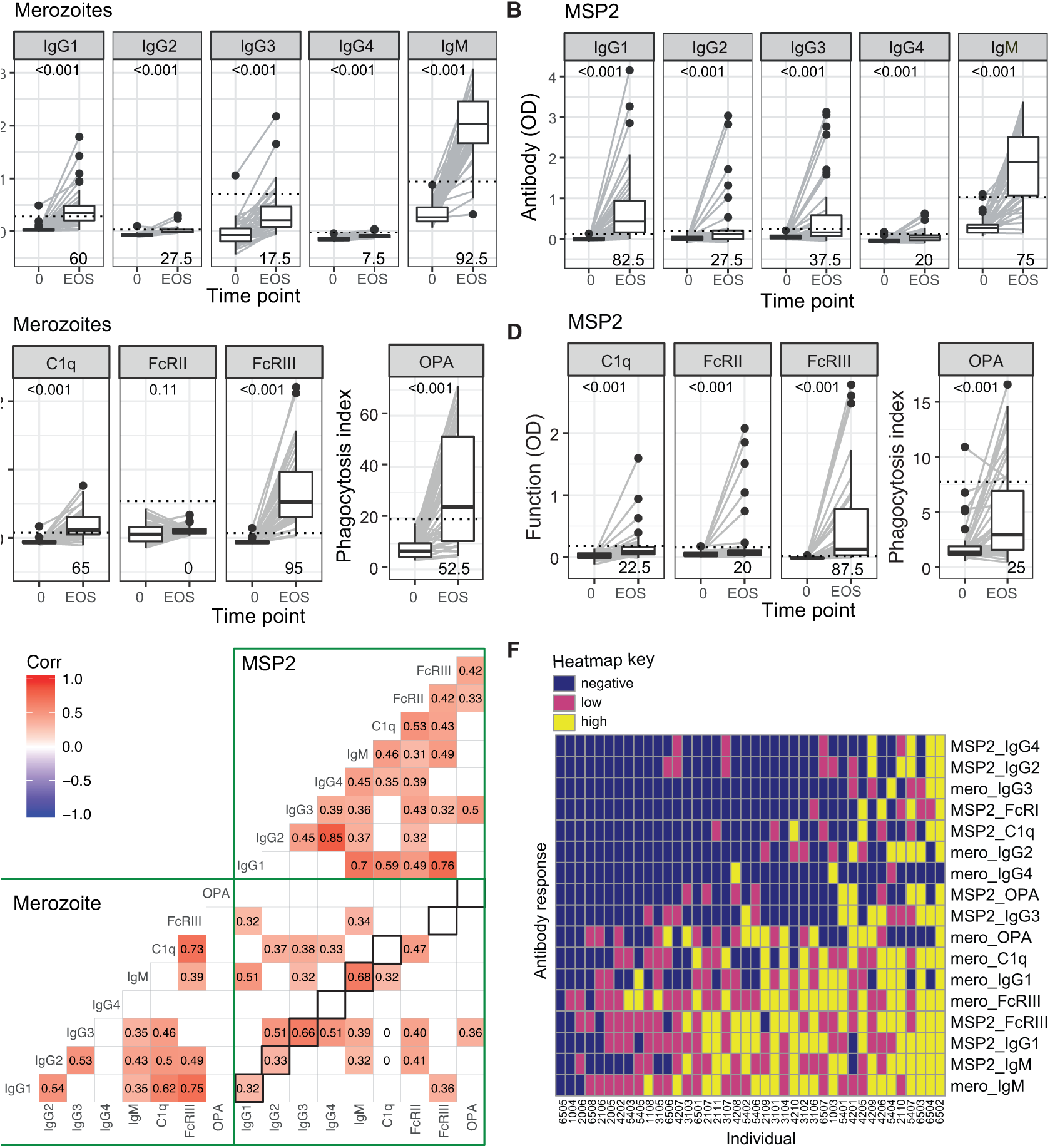
Antibodies to merozoite and MSP2 were induced following IBSM. **A-D)** IgG subclasses and IgM responses and functional capacity to fix complement (C1q), binding of dimeric Fc_γ_RIIa or Fc_γ_RIIIa (surrogates for IgG antibody capacity to cross-link cellular receptors) and to promote opsonic phagocytosis (OPA) to merozoites **(A/C)** and MSP2 **(B/D)** were assessed prior to inoculation and at end-of-study time points (EOS). Positive threshold for each responses is indicated by dotted line (calculated as mean + 3 SD of day 0 responses and the proportion of positive responders is indicated in the bottom right of each panel. Wilcoxon rank-sign t test is <0.001 for all comparisons of antibody responses at day 0 to EOS, except for merozoite Fc_γ_RIIα which was not significant p=0.11. **E)** Correlation matrix of all antibody responses to merozoites and recombinant MSP2. Only correlations <0.05 are indicated. Correlations between the same subclass/function of responses comparing merozoite and MSP2 responses are indicated in black squares. **F)** Heat map of composition of induced antibodies following categorization as negative (below positive threshold), or low/ high (based on below/above median value of all positive responses). Subjects are ranked in order of total breadth and magnitude of response.

### Antibody induction is associated with Th2-cTfH activation during peak infection

To investigate for any links between cTfH cell activation and antibody induction, we calculated an antibody score for each individual antibody response based on categorized responses (negative, low, high were given a score of 0, 1, 2 and then summed). This approach captured the combined breadth and magnitude of antibody responses regardless of antibody composition (Figure 6A). We found that the antibody score was positively associated with the increase of ICOS+ cTfH cells at day 8 (compared to d0), but not other time points (Figure 6B). The positive association between antibody score and ICOS+ cTfH cells at day 8 was restricted to the Th2-cTfH cell subset (Figure 6C). This relationship remained after removal of outliers (**Supplementary Figure 5**). Neither antibody score, nor individual antibody responses, were associated with parasite biomass during infection (**Supplementary Figure S6A/B)**, in contrast to previous reports *(48–50)*.

**Figure 6:**
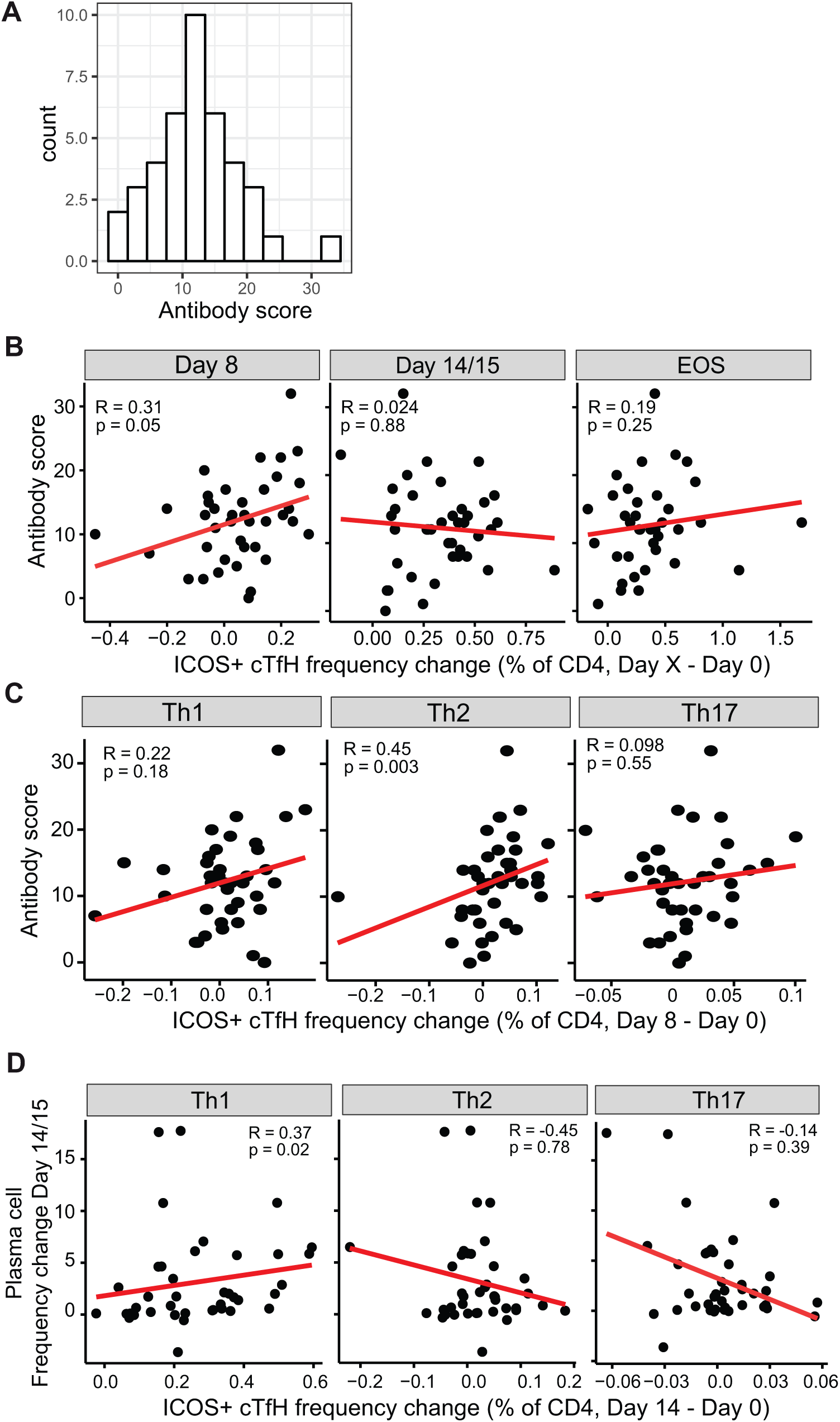
Th2-TfH cells are associated with antibody induction and Th1-TfH cells are associated with plasmacell expansion. **A.** To assess total breadth and magnitude of antibody response, responses were categorized as negative (0), or low (1) and high (2) responds and combined to calculate a total antibody score. **B.** Association between antibody score and the frequency change of ICOS+ cTfH cells compared to day 0 at each time point. **C.** Association between antibody score and the frequency change of each subset of ICOS+ cTfH cells at day 8. **D.** Association between plasma cell induction and ICOS+ cTfH cells at day 14/15 following infection. Spearman correlation and p are indicated. See also Supplementary Figure S5 & S6.

It has been hypothesised that short lived plasma cell differentiation may be driven by Th1-cTfH cells *(29)*, and Th1-cTfH cell activation is associated with plasma cell induction following influenza vaccination *(19)*. Consistent with this, there was a positive association between activated Th1-cTfH cells and plasma cell expansion at day 14/15 (Figure 6D); we have previously shown that plasma cells are induced at day 14/15 in experimental infections *(30)*. Plasma cell expansion was not associated with antibody responses at EOS (r=0.078, p=0.3).

## Discussion

Despite decades of research, the most advanced malaria vaccine, RTS,S has only ~30% efficacy in target populations. TfH cells are the key drivers of antibody induction, and optimized activation of TfH cells during vaccination may boost malaria vaccine efficacy *(7, 51)*. Here, we prospectively investigated the role of TfH activation in antibody responses to malaria infection during experimental *P. falciparum* infection in malaria-naïve adults. scRNAseq of cTfH during infection indicated that the highest transcriptional activation of multiple cTfH cell subtypes occurred during peak infection, at which point infection driven clonal expansion was evident. Subsequently, in a large cohort of individuals, we show that activation of Th2-cTfH cells occurs during peak infection and this activation is positively associated with antibody magnitude and functional activities. In contrast, Th1-cTfH cell activation occurs after anti-malaria drug treatment and was associated with plasma cell development, which we have shown has negative impacts on the acquisition of long lived and memory humoral responses *(30)*.

Understanding the drivers of functional antibody responses will assist in the development of vaccines with the highest protective efficacy. Here, using experimental malaria infection, a powerful infection model that allows for the longitudinal study of immune responses in previously naïve individuals free of confounding environmental factors commonly seen in cohort studies, we find that early Th2-cTfH cell activation is associated with antibody induction during malaria. These findings open avenues to identify strategies for generating more potent malaria vaccines. Indeed, targeted TfH cell activation is beginning to be explored as an avenue to improve malaria vaccine efficacy, and antigen specific cTfH has been identified as one predictor of RTS,S efficacy *(8)*. Strategies such as using a glycopyranosyl lipid adjuvant-stable emulsion have been shown to increase the quantity of TfH cell activation (without changing the composition of the response), resulting in increased magnitude and longevity of induced anti-malarial antibodies in humans *(51)*. However, the co-administration of viral vectored vaccines with RTS,S skewed the cTfH cell response towards Th1-cTfH cell activation to the detriment of functional antibody induction and vaccine efficacy *(52)*. These findings are consistent with our data of the specific importance of Th2-cTfH cells in antibody activation, in contrast to the potential for Th1-cTfH cell responses to hamper immune development, and highlight the importance of targeting specific TfH cell subsets to improve vaccine efficacy.

Antibody induction was associated with early cTfH activation during peak infection, a finding consistent with our scRNAseq analysis showing highest activation of cTfH and clonal expansion of putative parasite specific cells at this time point. This is also in agreement with previous studies indicating a key role of early activation of cTfH in providing B cell help during initial phases of GC formation following experimental HIV vaccinations *(53)*. The specific association between Th2-cTfH cell activation and antibody acquisition is in line with known properties of Th2-cTfH cells which have the greatest capacity to activate naïve B cells *(10)* and are transcriptionally most similar to GC TfH cells *(9)*. Here we confirmed by scRNAseq that amongst circulating CXCR5+ CD4 T cells, Th2-like cells have highest GC TfH signatures. ScRNAseq allows for the unprecedented ability to characterise the heterogeneity of cell populations. We have recently applied these techniques to study CD4 T cell development during murine malaria models *(54)*, and now for the first time, we have applied such approaches to cTfH cells during experimental human malaria. While previous studies have suggested that cTfH cells may exist in 5-7 distinct clusters based on surface markers and cytokine production *(55, 56)*, our scRNAseq analysis suggests that the current phenotypical strategy of defining cTfH cell subsets captures the majority of variation within cTfH cells most similar to GC populations. We identified Th2-, Th1-, Th17- and TfReg clusters with characteristics shared with these previously identified populations, with no other major clusters with high GC gene transcriptional signatures nor clear transcriptional profiles identified. Of note, scRNAseq analysis resulted in finding that Th2-cTfH cells have relatively lower expression of CXCR4. CXCR4 drives cellular sub-location within the GC light and dark zones, with low CXCR4-expressing cells locating to the light zone and high CXCR4-expressing cells locating to dark zone *(35)*. In the light zone, B cells are selected based on BCR affinity, while in the dark zone B cells rapidly proliferate and undergo somatic hyper mutation. As such, both expression of GC TfH cell signatures, which is highest in Th2-like cTfH cells, and relative CXCR4 expression may contribute to TfH cell subset function.

During experimental *P. falciparum* infection, the dynamics of Th2- and Th1-cTfH cell activation appear to be distinct. Th2-cTfH cell activation was detected early at peak infection, while Th1-cTfH cell phenotypic activation was not seen until one week after parasite treatment. Data presented here and previously *(12)* suggest that Th2- and Th1-cTfH cells are clonally divergent, consistent with distinct development linages. Little is known regarding the specific induction of Th2-cTfH cells during infection, but transcriptional factor *Ets1* has been reported to be a negative regulator of Th2-TfH cells *(57)*. For Th1-TfH cell activation, preferential induction during *Plasmodium* infection is thought to be mediated by a parasite driven Type I IFN associated IFNγ production *(24, 58)*. We have previously shown that IFN production by CD4+ T cells peaks one week after treatment in human experimental infection *(59)*, possibly due to inflammatory responses following parasite death, and consistent with the peak of Th1-cTfH activation seen here. Studies by ourselves and others have indicated that excessive Type I IFN signaling during *Plasmodium* infection in mice can suppress ICOS-dependent antibody mediated immunity *(60)*, and induce regulatory response that restrict TfH induction *(59, 61)*. However, Type I interferon signaling is also clearly an important activator of TfH cells, with Type I IFN signaling signature genes increased in cTfH cells during peak infection across multiple TfH cell clusters identified by scRNAseq, including within Th2-like cells. Further studies are required to identify the factors that control the activation and development of Th2-cTfH cells distinct from Th1-cTfH cells.

In contrast to the role of Th2-cTfH cells in inducing antibodies, during IBSM Th1-cTfH cell activation was associated with plasma cells. We have recently shown that plasma cells induced during *Plasmodium* infection are metabolically hyperactive and act as a nutrient sink that limits the generation of protective GC responses and suppresses long term humoral immunity *(30)*. Here, we also show that Th1-cTfH cells are the TfH subset that express granzyme B. Expression of granzyme B in TfH cells has been previously identified in mice *(62)* and humans *(63, 64)* and has been implicated in TfH cell-mediated lysis of B cells in group A streptococcal infection *(64)*. Th1-cTfH cells induced during malaria have also been implicated in the induction of atypical memory B cell responses *(26)*, which expressed FCRL5. These B cells have previously been considered as ‘exhausted’ memory B cells which contribute to the slow acquisition of protective anti-malarial humoral responses in naturally exposed populations *(27, 28, 65)*. It has recently been shown that FCLR5+ B cells rapidly give rise to antibody producing B cells during secondary infection, and that these are present in memory responses induced by vaccination *(66)*. Together these data suggest that the induction of Th1-cTfH cells during malaria may have multiple roles in negatively regulating GC responses. Indeed previous studies have shown that cTfH activation during *P. falciparum* malaria in children is restricted to Th1-cTfH cells *(24)*, and it has been hypothesized that this Th1-cTfH bias underlies the slow acquisition of malaria immunity in children *(67)*. Further studies are required to understand the similarities and differences between cTfH cell activation and antibody responses in experimental malaria in naïve adults, compared to naturally exposure with *Plasmodium*, which in most settings occurs predominantly in children.

In conclusion, our findings demonstrate for the first time a link between a specific subset of TfH cells that is required for the induction of robust and functional anti-malarial antibodies. These studies improve our understanding of the dynamics of TfH cell activation during *Plasmodium* infection in humans and have led to the identification of Th2-cTfH cells as specific TfH subsets driving anti-malarial antibody production. Due to the key role of functional antibodies in vaccine induced and naturally acquired anti-malarial protective immunity, data for the first time identifies Th2-cTfH as a cell subset that can be targeted to improve anti-malarial vaccine efficacy.

## Methods

### Study approval

Written informed consent was obtained from all participants or, in the case of children <18 years of age, from their parents or guardians prior to inclusion in studies. Ethics approval for the use of human samples in the relevant studies was obtained from the Alfred Human Research and Ethics Committee for the Burnet Institute (#225/19), and from the Human Research and Ethics Committee of the QIMR-Berghofer Institute of Medical Research Institute (P1479 and P3445). Ethical approval for the clinical trials from which the IBSM study samples were collected was likewise gained from the Ethics Committee of the QIMR-Berghofer Medical Research Institute.

### Study cohorts

Induced Blood Stage Malaria inoculum preparation, volunteer recruitment, infection, monitoring and treatment were performed as previously described *(46)*. In brief, healthy malaria-naive individuals underwent induced blood-stage malaria inoculation with 2800 viable *P. falciparum* 3D7-parasitized RBCs, and peripheral parasitemia was measured at least daily by qPCR as described previously *(68)*. Participants were treated with antimalarial drugs at day 8 of infection when parasitemia reaches approximately 20,000 parasites/ml. Blood samples from 40 volunteers (from 4 studies across 6 independent cohorts) were collected prior to infection (day 0), at peak infection (day 8) and 14 or 15 and 27-36 days (end of study, EOS) after inoculation (in analyses these time points are grouped as 0, 8, 14/15 and EOS). Plasma was collected from lithium heparin whole-blood samples according to standard procedures, snap frozen in dry ice and stored at −70°C. Participants were healthy malaria naïve adults (n=40 (90% male), age 25.5 [21.25-31], median[IQR]) with no prior exposure to malaria or residence in malaria-endemic regions. Clinical trials were registered at ClinicalTrials.gov NCT02867059 *(69)*, NCT02783833 *(70)*, NCT02431637 *(71)*, NCT02431650 *(71)*. PBMC and plasma from healthy non-infected controls was collected by the same processes (n=8 (38% male), age 40[33-56] median[IQR], for samples in **Supplementary Figure S1**).

### Calculation of parasite density during infection

Area under the curve (AUC) were calculated using the trapezoidal method on serial log_10_ transformed parasites/mL data from 4 days p.i. to each of the three defined timepoints (8, 14/15, and EOS as described previously *(30)*. Equation 1 below describes the calculation, with *t*_*i*_ being each time point sampled, *P*_*i*_ being the log_10_ parasites/mL at that time, and *T*_*N*_ being either 8 p.i., 14/15 p.i., or EOS.

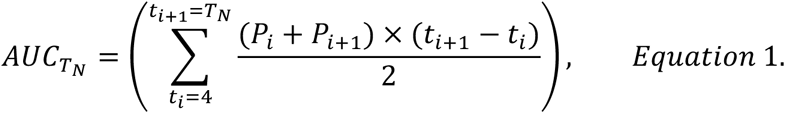

Samples where parasitaemia was not detected were substituted with 0 on the log_10_ scale. The samples collected between the 4 defined timepoints (ranging from daily to twice daily before treatment, ranging from daily to every two hours after treatment, and ranging from every four days to daily between timepoint 14/15 and EOS) were used in the calculation of AUC but not in any other analyses.

### Flow cytometry

200μl of whole blood or thawed PBMCs were stained with antibodies as outlined in **Supplementary Table S2.** For whole blood, RBCs were lysed with FACS lysing solution (BD) and resuspended in 2% FCS/PBS, and acquired on the BD LSRFortessa 5 laser cytometer (BD Biosciences). For PBMCs staining, cells were permeabilized (Fix Perm, Ebioscience) and stained with intracellular FoxP3 (206D) and Granzyme B (QA16A02), samples were re-suspended in 2% FCS/PBS and acquired on the Aurora (Cytek Biosciences, USA). These data were analyzed using FlowJo software (Tree Star, San Carlos, CA, USA) and all antibodies were purchased from BD Biosciences or Biolegend (**Supplementary Table S2**).

### Single cell RNA sequencing

Frozen human PBMC samples were thawed at 37°C and then stained for flow cytometry assessment. Cells were stained with LIVE/DEAD™ Fixable Near-IR Dead Cell Stain Kit (Thermo Fisher Scientific), CD14 – Qd605 (TüK4; Thermo Fisher Scientific), CD3 – PE-CF594 (UCHT1; BD Biosciences), CD4 – BV510 (OKT4; Biolegend), CD45RA – BB535 (HI100; BD Biosciences) and CXCR5 – BV421 (RF8B2; BD Biosciences). cTfH CD4^+^ T cells were sorted as live singlet CD14^−^, CD3^+^ CD4^+^, CD45RA^−^ CXCR5^+^. The sorted cells were loaded onto a Chromium controller (10x Genomics) for generation of gel bead-in-emulsions. Approximately 8,700 cells were loaded per channel and subsequent scRNAseq library preparation was performed using the Single Cell 5’ Reagent Kit (10x Genomics), according to the manufacturers’ protocol. scRNAseq libraries were pooled and sequenced on an Illumina NextSeq 550.

Raw sequencing data was processed using Cell Ranger version 2.1.0 with 10x human genome (hg19) 1.2.0 release as a reference. Data was filtered to remove cells expressing less than 200 genes and more than 4000 genes, and cells with greater than 11% mitochondrial content. Only genes expressed in 5 or more cells were considered. For downstream computational analyses, Seurat version 3.0.3 was used. SCTransform was applied on the high quality data to perform normalisation using regularised negative binomial regression as described*(72)*. Datasets from each time point were integrated as described *(73)*. Briefly, total of 2000 features were identified from “SelectIntegrationFeatures” and then integration anchors identified from “FindIntegrationAnchors” were used to integrate the datasets. Top 30 principal components were used for UMAP dimensionality reduction. Cell clusters were identified using the Louvain algorithm implemented in “FindClusters” command. Differential gene expression analysis was performed with each cluster against all other clusters using a Wilcoxon Rank Sum test implemented in “FindConservedMarkers” command. All genes with False Discovery Rate (FDR) below 0.01 were considered significant. For TCR analysis, only high-confident clones that had a single TRA and TRB chain were analysed. Cells were considered clones if TRA/TRB were concordant. Clonal TCR sequence relationships within and between timepoints were visualized using R package Circlize (Version 0.4.9) *(74)*.

### Antibodies to merozoites and recombinant MSP2

The level of antibody isotypes targeting merozoites were measured by standard ELISA methods as previously described (17). 96-well flat bottom MaxiSorp® plates (Nunc) were coated with 50 µl of merozoites (2.5 × 10^5^ merozoites/ml) or 50 µl of 0.5 µg/ml MSP2 recombinant antigen *(46)* in PBS overnight at 4°C. Plates were blocked with 150µl of 10% skim milk in PBS for merozoites or 1% casein in PBS (Sigma-Aldrich) for MSP2 for 2 hours at 37°C. Human serum samples were diluted in 0.1% casein in PBS and incubated for 2 hours at RT. For IgG detection, plates were incubated with goat polyclonal anti-human IgG HRP-conjugate (1/1000; Thermo Fisher Scientific) for 1 hour at RT. For detection of IgG subclasses and IgM, plates were incubated with a mouse anti-human IgG1 (clone HP6069), mouse anti-human IgG3 (HP6050) or mouse anti-human IgM (clone HP6083) at 1/1000 (Thermo Fisher Scientific) for 1 hour at RT. This was followed by detection with a goat polyclonal anti-mouse IgG HRP-conjugate (1/1000; Millipore). For all ELISAs, plates were washed three times with PBS-Tween 0.05% between antibody incubation steps. Serum dilution used for merozoites was 1/100 for IgG, 1/250 for IgG subclasses and IgM, 1/100 for C1q and FcR. Serum dilution used for MSP2 was 1/100 for IgG, 1/250 for IgG subclasses and IgM, 1/100 for C1q and 1/50 for Fc_γ_R. For detection of complement fixing antibodies, following incubation with human sera, plates were incubated with purified C1q (10μg/ml; Millipore) as a complement source, for 30 min at RT. C1q fixation was detected with rabbit anti-C1q antibodies (1/2000; in-house) and a goat anti-rabbit-HRP (1/2500; Millipore). TMB liquid substrate (Life Technologies) was added for 1 hour at RT and the reaction was stopped using 1M sulfuric acid. The optical density (OD) was read at 450 nm. For ELISA using isolated merozoites, washes were done with PBS without Tween to prevent parasite lysis. Standardization of the assays was achieved using positive control plasma pools on each plate. Background values (wells with no plasma) were subtracted from all values of other wells, and positivity was determined as the mean plus 3 standard deviations of the OD from naïve plasma samples at day 0 prior to inoculation. To assess how induced responses compared to the magnitude of antibodies in naturally exposed immune adults, antibody responses were normalized to the level of antibodies in a sera pool of immune adults from Papua New Guinea (PNG), an area of high malaria endemicity. To investigate the relationship between responses, the positive cut-off threshold was subtracted from all responses (as measured by OD), and then normalized to positive control (pool of hyperimmune PNG adults).

### Fcγ receptor-binding assay

A standard ELISA protocol was modified to measure the level of antibody-mediated FcγR binding *(75, 76)*. 100uL of recombinant protein at 0.5 ug/ml was coated on Maxisorp™ plates (Nunc) and incubated overnight at 4°C, followed by 3 washes with PBS-Tween. The plates were then blocked with 200μL of 1% BSA in PBS (PBS-BSA) at 37°C for 2 hours followed by 3 washes with PBS-Tween. Serum samples were diluted at 1:100 in PBS-BSA and 100μL of each sample was added to the ELISA plates in duplicate and incubated at room temperature for 2 hours, followed by 3 washes in PBS-Tween. 100ul of biotin-conjugated rsFcγRIIa H131 or rsFcγRIIIa V158 ectodomain dimer (0.2ug/ml) was added to each well and incubated at 37°C for 1 hour followed by 3 washes with PBS-Tween. This was followed by a secondary horseradish peroxidase (HRP)-conjugated streptavidin antibody (1:10,000) in PBS-BSA at 37°C for 1 hour followed by 3 washes with PBS-Tween. Finally, 50 μL of TMB liquid substrate was added for 20 minutes to measure enzymatic reactivity. The reaction was stopped with 50 μL of 1M sulfuric acid solution. The level of binding was measured as optical density at 450 nm. Pooled human IgG from malaria-exposed Kenyan adults (1/100) and rabbit polyclonal IgG to PfCSP (1:500) was used as positive controls. Individual sera from naive Melbourne adults (1/100) were used as negative controls.

### Coating fluorescent latex beads with antigen

Amine-modified fluorescent latex beads (Sigma) were washed twice with 400ul of PBS and centrifuged at 3000g for 3 min *(76)*. 400uL of 8% glutaraldehyde (diluted in PBS) was added to the beads and incubated on a roller overnight at 4°C. After washing with PBS, 1mg/ml of recombinant MSP2 was added to the mixture and incubated on a vortex for 4 hours. Subsequently, the mixture was centrifuged, and the pellet was collected as the bound protein fraction. 200ul of ethanolamine was added to the pellet to quench amine groups and incubated for 30 min on the vortex. The pellet was subsequently washed in PBS and blocked with 1% BSA overnight at 4°C. The antigen-coated beads were stored 4°C in the presence of 0.1% SDS and 0.02% sodium azide.

### Opsonic phagocytosis of antigen-coated beads with monocytes

The density of latex beads coated with recombinant MSP2 was adjusted to 5×10^7^ beads/ml and opsonised with serum samples (1/10 dilution) for 1 hour *(47, 76)*. Samples were washed thrice with RPMI-1640 before co-incubation with THP-1 monocytes for phagocytosis. Phagocytosis was allowed to occur for 20 min at 37°C and samples were subsequently washed with FACS buffer at 300g for 4 minutes. The proportion of THP-1 cells containing fluorescent-positive beads was evaluated by flow cytometry (FACS CantoII, BD Biosciences) and analysed using FlowJo software.

### Data analysis

Data analysis was performed in RStudio 1.1.456 or GraphPad Prism 7. Paired data of immune responses between time points was analysed by Wilcoxon paired t-test. Correlations between responses were analysed by Spearman’s correlation.

## Author Contributions

MJB, CE, JGB, AH designed research studies

JAC, FdLR, JRL, JE, ASN conducted experiments

JAC, FdLR, JRL, JE, PM, AMP, ASN, MJB, LW analysed data

FHA, JSM, JGB, BDW, PMH provided essential reagents

MJB, JAC,CE, AH wrote manuscript with feedback and approval from all other authors

### Acknowledgements

RBC and human serum used for parasite culture and naïve control sera were provided by the Australian Red Cross Blood Bank (Melbourne). We acknowledge Robin Anders for providing the recombinant MSP2 antigen. We thank the participants involved in the malaria volunteer infection studies, Q-Pharm staff, and Medicine for Malaria Venture for funding these studies.

