## Supplementary Figures for "Th2-like T-follicular helper cells promote functional antibody production during *Plasmodium falciparum* infection"

**Supplementary Material**

**Tables**

**Supplementary Table S1: Conserved cluster marker genes**

Attached file

**Supplementary Table S2: Flow cytometry antibodies for cellular studies**

| Panel | Antigen | Fluorochrome | Clone | Supplier |
|---|---|---|---|---|
| VIS whole blood | CD20 | BUV395 | 2H7 | BD Biosciences |
| | CXCR5 | BV421 | RF8B2 | BD Biosciences |
| | CD4 | VH500 | RPA-T4 | BD Biosciences |
| | CCR6 | BV650 | 11A9 | BD Biosciences |
| | CD38 | BV785 | HIT2 | BD Biosciences |
| | CXCR3 | APC | 1C6 | BD Biosciences |
| | CD27 | AF700 | M-T271 | BD Biosciences |
| | CD8 | APC-Cy7 | SK1 | BD Biosciences |
| | CD19 | FITC | HIB19 | BD Biosciences |
| | CD45 | PerCP-Cy5.5 | 2D1 | BD Biosciences |
| | ICOS | PE | DX29 | BD Biosciences |
| | CD3 | PE-CF594 | UCHT1 | BD Biosciences |
| | PD1 | PE-Cy7 | EH12.1 | BD Biosciences |
| Control PBMCs | Foxp3 | BV421 | 206D | Biolegend |
| | CXCR3 | Pacific Blue | G025H7 | Biolegend |
| | CXCR4 | BV510 | 12G5 | Biolegend |
| | CXCR5 | BV711 | J252D4 | Biolegend |
| | CCR6 | BV650 | 11A9 | BD Biosciences |
| | CD127 | BV570 | A019D5 | Biolegend |

| | | | | |
|---|---|---|---|---|
| | CD3 | FITC | SK7 | Biolegend |
| | CD4 | PerCP-Cy5.5 | OKT4 | Biolegend |
| | PD-1 | PE-Cy7 | EH12.1 | BD Biosciences |
| | Granzyme B | APC | QA16A02 | Biolegend |
| | CD25 | AF700 | 2A3 | BD Biosciences |
| | ICOS | APC-Cy7 | C398.4A | Biolegend |
| | Live/dead | NIR | | Biolegend |

**Supplementary Figure S1**

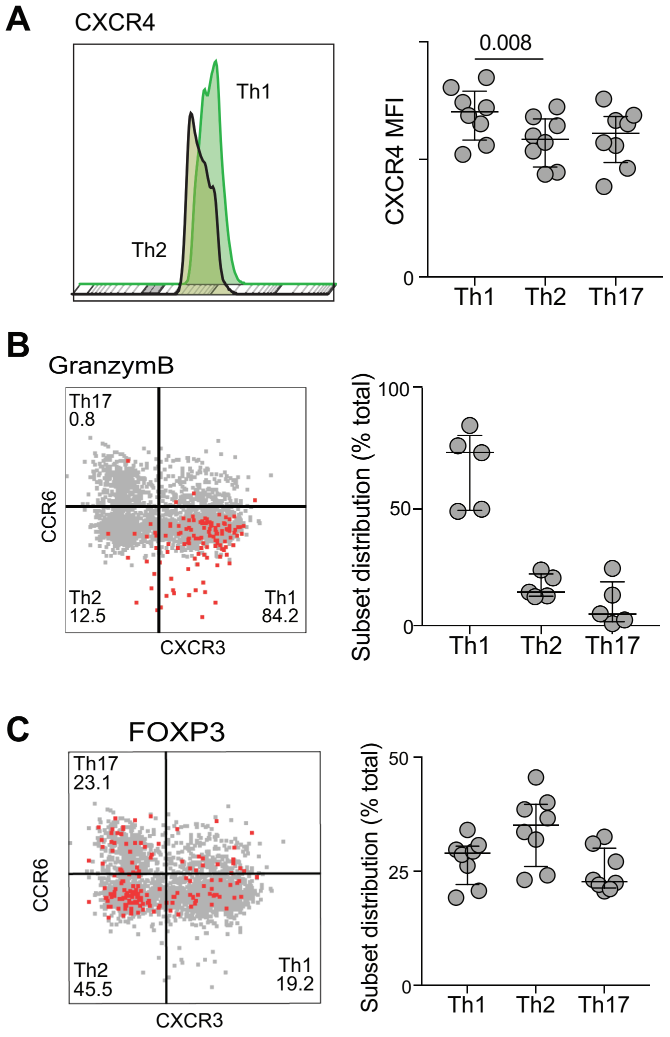

**Supplementary Figure S1: Flow cytometry analysis of cluster features.**

CXCR4 (**A**), Granzyme B **(B)**, and FoxP3 **(C)** expression was assessed within cTfH subsets in five-eight healthy blood donors. Left panels are representative plots of a single donor. Right panels are combined data.

**Supplementary Figure S2**

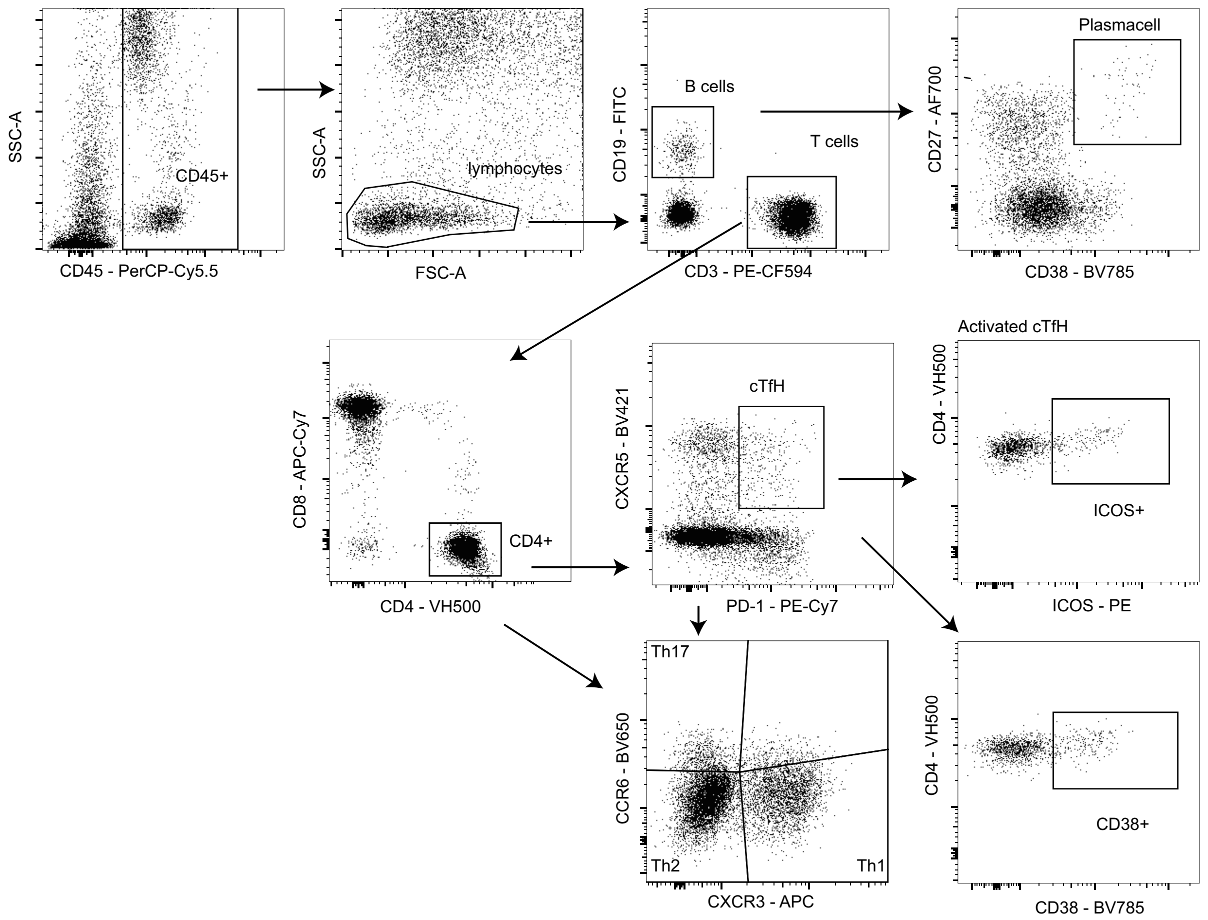

**Supplementary Figure S2: Gating strategy for TfH and plasma cells**

Gating strategy to identify cTfH, including activation and subsets, and antibody secreting cells (ASCs). Whole blood was stained and analysed by flow cytometry, CD45+ lymphocytes were gated as T-cells on CD3 or B cells on CD19. CD4 T cells were gated as CD4+CD8- and cTfH analysed based on PD1+CXCR5+ cells. cTfH subsets were analysed based on CXCR3 and CCR6 staining into Th1 (CXCR3+CCR6-), Th17 (CXC3-CCR6+) and Th2 (CXCR3-CCR6-) subsets. Activation was gated as ICOS+ and CD38+. Plasma cells were gated as CD38+CD27+ B cells.

**Supplementary Figure S3**

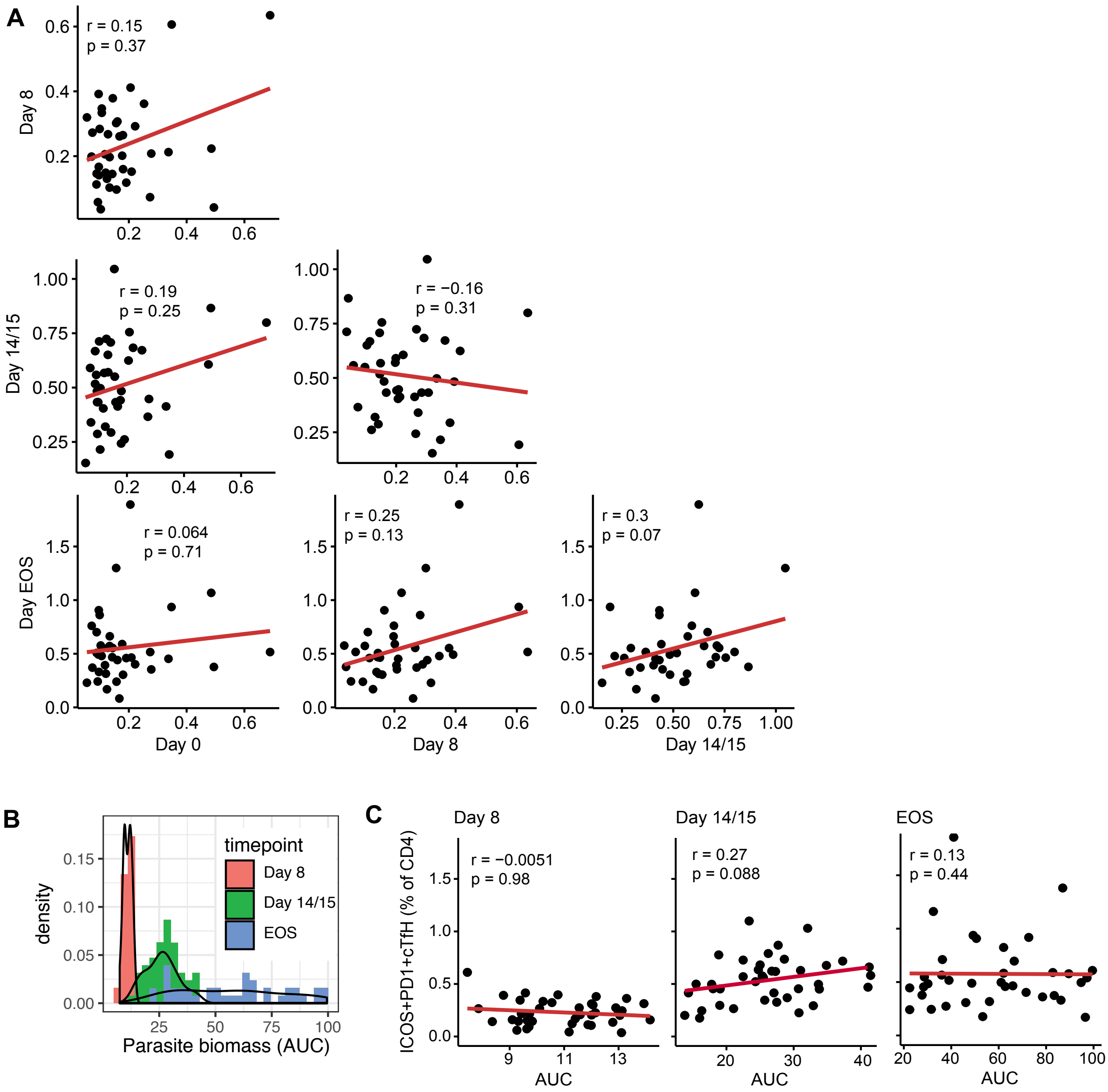

**Supplementary Figure S3 Correlations between ICOS+ cTfH cells and parasite biomass**

**A**. Correlations between ICOS+ cTfH (PD1+CXCR5+, data is % of CD4 T cells) at each time point (day 0, 8, 14/15 and end-of-study). Spearman's correlations coefficients and *p* values indicated.

**B**. Parasite biomass was calculated as Area Under the growth Curve (AUC) on the $\log_{10}$ scale from inoculation to day 8 (day of treatment), day 14/15 and end-of-study time points from qPCR data.

**C**. Correlations between ICOS+ cTfH and parasite density (area under curve, AUC) at each time point.

**Supplementary Figure S4**

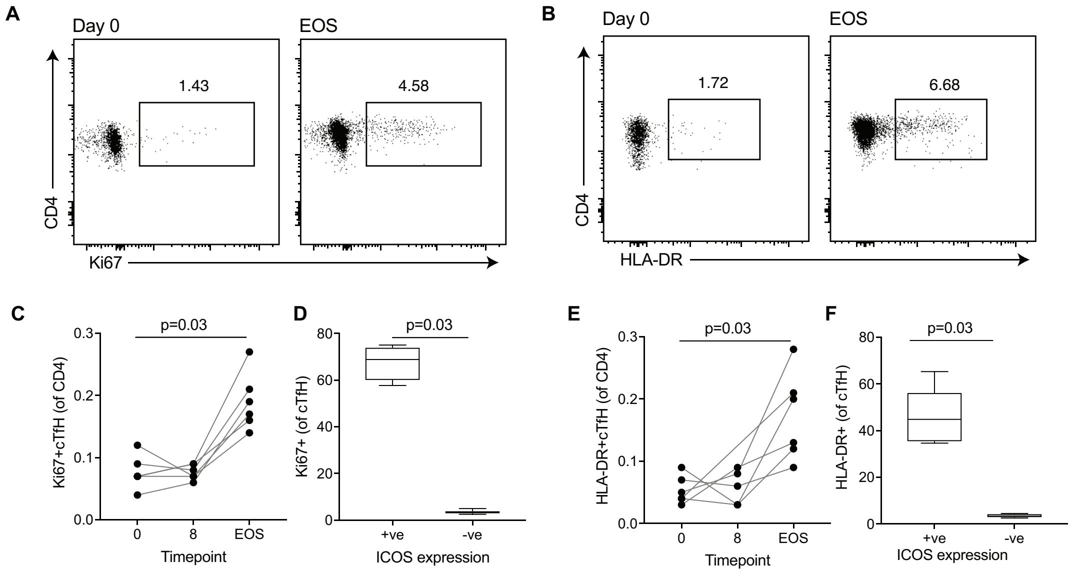

**Supplementary Figure S4: Ki67 and HLA-DR expression on cTfH**

PBMC samples from a subset of 6 participants were assessed for Ki67 **(A)** and HLA-DR **(B)** expression on cTfH (PD1+CXCR5+) at days 0, 8 and EOS time-points. The frequencies of Ki67+ **(C)** and HLA-DR+ **(E)** PD1+cTfH increased during experimental infection. Expression of Ki67 and HLA-DR was higher on ICOS+ than ICOS- cTfH subsets **(D**, **F)**.

**Supplementary Figure S5**

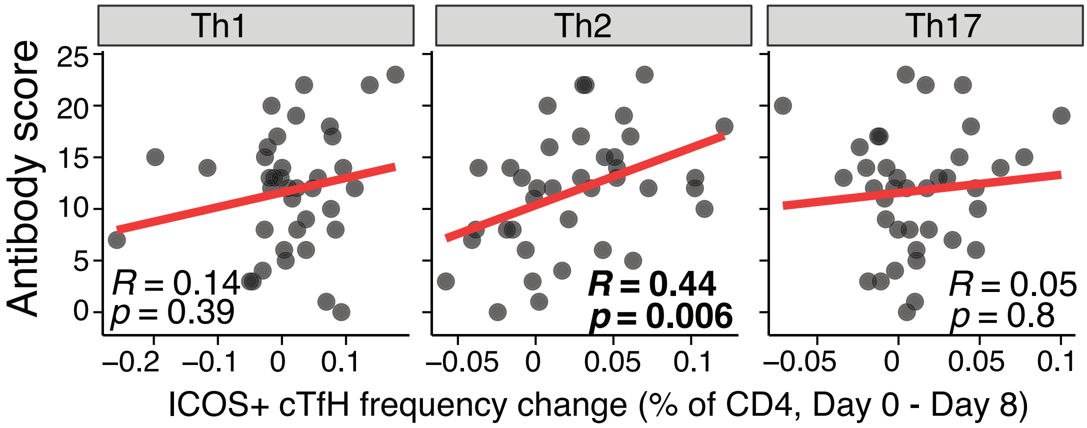

**Supplementary Figure S5: Th2-TfH cells are associated with antibody induction.**

Association between antibody score and the frequency change of each subset of ICOS+ cTfH cells at day 8 following removal of two individuals with outlier antibody scores or ICOS+ cTfH frequency changes.

### Supplementary Figure 6

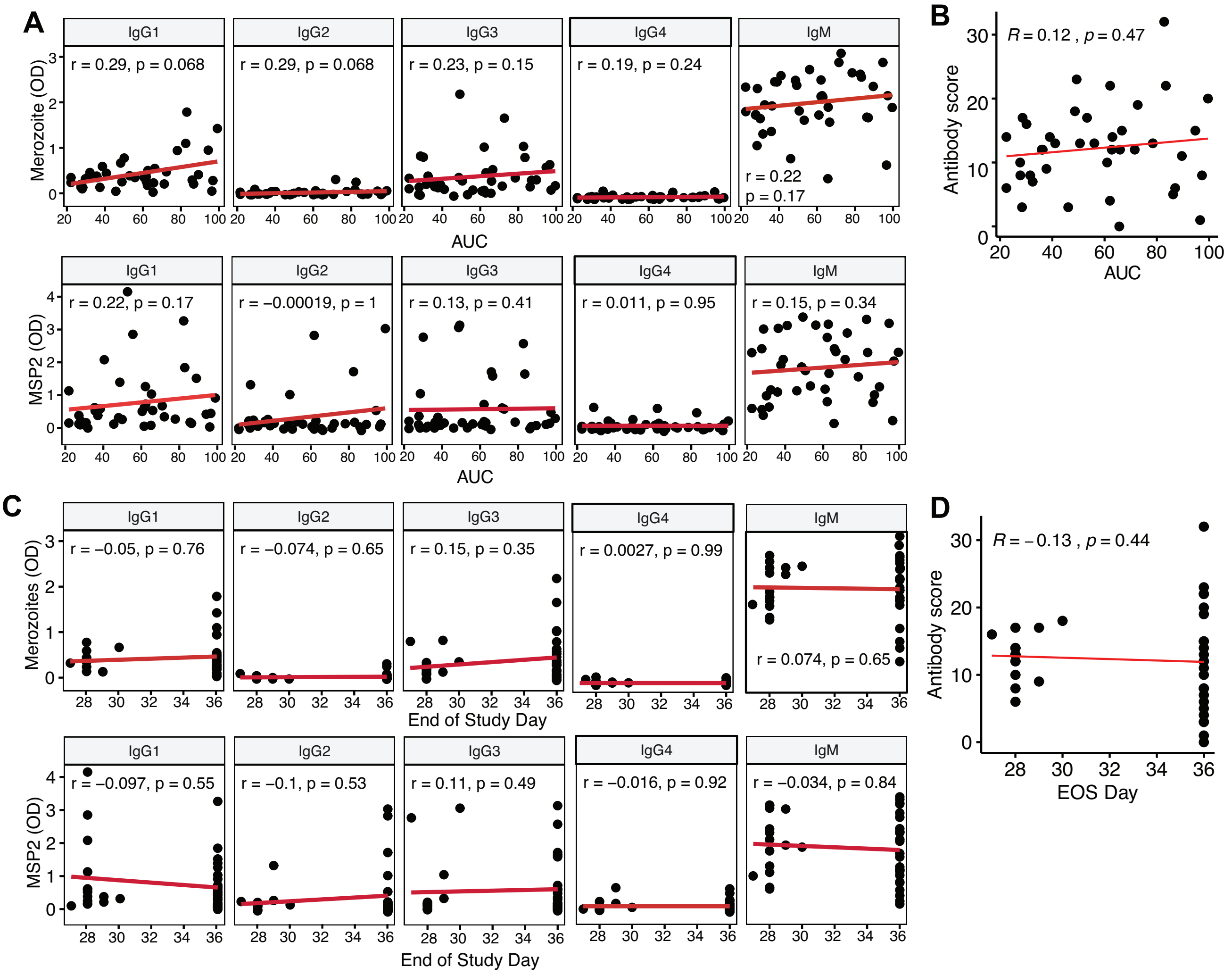

**Supplementary Figure S5 Relationship between antibodies and parasite burden**

**A**. Correlation between individual antibody responses to merozoites and MSP2 with parasite biomass as measured by area under curve at EOS time point.

**B**. Correlation between total antibody score with parasite biomass at EOS time point.

**C**. Correlation between individual antibody responses to merozoites and MSP2 and EOS day.

**D**. Correlation between total antibody score and EOS day.

Correlations were evaluated using Spearman's rho.
